# Deciphering chromosome fusion in *D. miranda’s* neo-sex chromosome through single-copy and repetitive oligo probes

**DOI:** 10.64898/2026.01.20.700498

**Authors:** Henry Bonilla Bruno, Isabela Almeida, Maria Dulcetti Vibranovski

## Abstract

*Drosophila miranda* is considered an excellent model for studying sex chromosome evolution due to its neo-sex chromosomes, which originated from fusions between autosomes and sex chromosomes. In this study, we took advantage of the latest genome assembly of *D. miranda* to design the first oligo probe libraries targeting neo-sex chromosomes, covering X and Y-linked regions with times ranging from ∼1.5 to 60 million years. These libraries, which include both single-copy and repetitive oligos, were generated by integrating the OligoY approach to the conventional OligoMiner pipeline and validated through fluorescence *in situ* hybridization (FISH). We optimized oligo density and spacing parameters to predict consistent and effective chromosome painting. Beyond tool improvement, our mapping of the three largest unplaced Y-linked scaffolds in *D. miranda* reveals a complex evolutionary mechanism driving the current structure of the Y chromosome, including chromosomal translocation, centromere loss, and inversions. This work provides essential tools for sex chromosome identification via probe labeling and offers a foundation for exploring the spatial and evolutionary dynamics of sex chromosomes across different cell types.

**Author summary:** While previous studies have focused on using single-copy oligonucleotides for chromosome painting, these oligos have limited effectiveness in targeting repetitive regions such as ribosomal DNA, pericentromeres, and mainly Y chromosomes. In this study, we integrated the OligoMiner and OligoY pipelines to design highly specific oligonucleotide libraries capable of targeting both single-copy and repetitive regions in any chromosome, enabling comprehensive painting of autosome and sex chromosomes. Using *Drosophila miranda* neo-sex chromosomes as a model, we validated the specificity of our oligo libraries through fluorescence in situ hybridization (FISH). Our results demonstrate that it is possible to achieve successful chromosome painting of sex chromosomes ranging from 1.5 to 60 million years old by combining single-copy and repetitive oligos, without compromising specificity. Notably, we painted the neo-Y chromosome of *D. miranda* and proposed a hypothesis to give rise to its current structure. This approach provides a powerful tool for studying chromosome evolution and organization, particularly in complex and repetitive genomic regions.

## Introduction

*Drosophila miranda*, part of the pseudoobscura subgroup, is distinct for having undergone two chromosomal fusion events involving sex chromosomes and autosomes (Steinemann M., 1982). These events led to the formation of neo-sex chromosomes, offering a unique platform to study evolutionary stages of X and Y chromosomes within one genome (Zhou et al., 2012; Zhou et al., 2013). Originally, the Muller-D element fused with Muller-A (15 Ma ago), forming the XL/XR chromosome and evolving into an X-like structure (Alekseyenko et al., 2013). Its homologous counterpart, the Muller-D element, underwent extensive degeneration and is suggested to represent the heterochromatic Y chromosome (referred to as YD) of *D. pseudoobscura* (Carvalho and Clark 2005) (Fig 1A). Later in the evolution of *D. miranda*, a second fusion ∼1.5 Ma combined the Muller-C element with YD, giving rise to the neo-sex chromosomes (neo-X and neo-Y). As a result, *D. miranda* bears distinct XL/XR (Muller-A and D) and Neo-X (Muller-C) chromosomes, alongside a YD/Neo-Y (YD/Muller-C) chromosome (Fig 1A).

**Fig 1.**
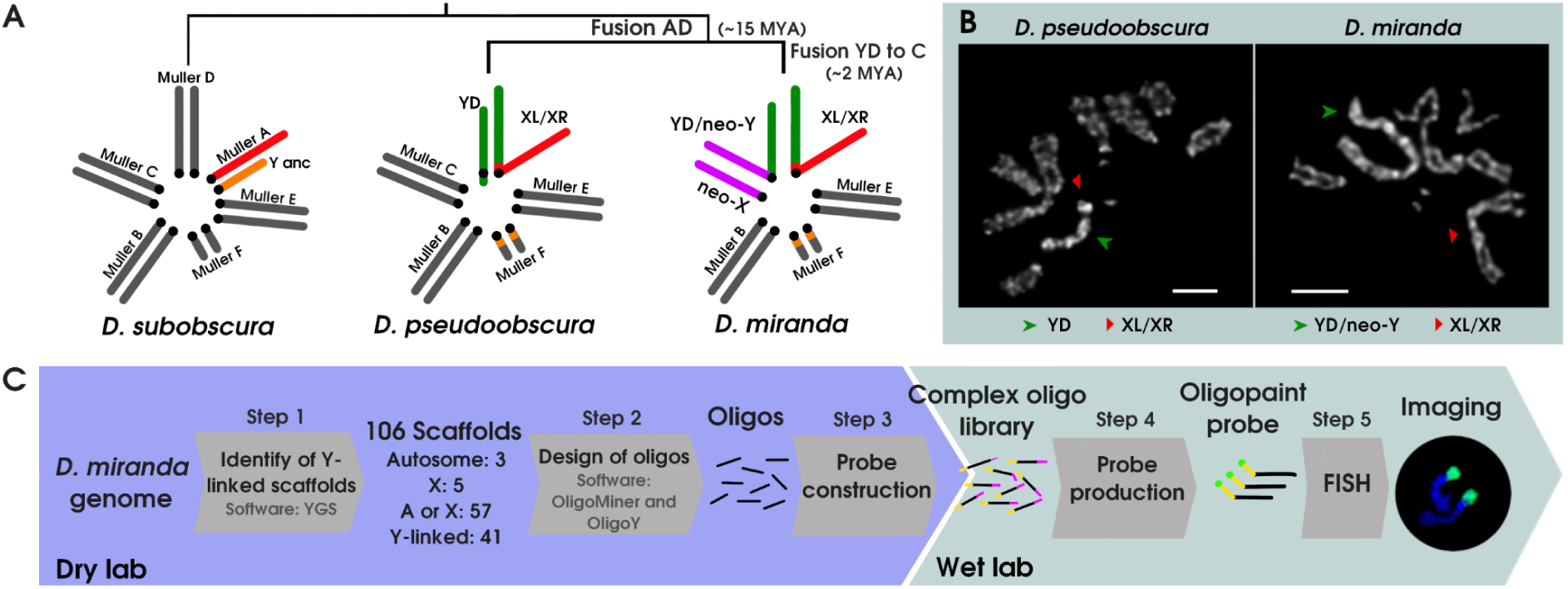
Evolution of Neo-Sex Chromosomes and Karyotype Diversity in the *Drosophila pseudoobscura* Subgroup, Along with Our OligoPaint Design Workflow. **A)** Evolution of the karyotype within the *pseudoobscura* subgroup leading to the formation of neo-sex chromosomes. Autosome Muller elements are presented in gray, while sexual Muller elements are shown in color. Muller A (red) represents the ancestral X. The fusion event between Muller A and D (green) is common to both *D. pseudoobscura* and *D. miranda*, resulting in XL/XR. However, the fusion between Muller C (magenta) and YD is specific to *D. miranda*, giving rise to the neo-X and YD/neo-Y chromosomes. Genes of the ancestral Y (orange) are located in the dot (chr 5) (Mahajan S. et al., 2018). **B)** Male metaphase of *D. pseudoobscura* (2n=10) and *D. miranda* (2n=9). **C)** Schematic overview illustrating the sequential steps to design and produce oligopaints.

Comparative analysis of homology blocks between the Neo-X and YD/Neo-Y chromosomes, in the most recent and reference genome of *D. miranda,* revealed that the YD/Neo-Y chromosome consists of a long and short arm (Mahajan et al., 2018). The long arm is believed to represent the neo-Y, while the short arm is thought to have originated from the YD of *D. pseudoobscura*. However, cytogenetic observations indicate that the Y chromosome is actually metacentric (Steinemann 1982) (Fig 1B). Thus, chromosome-specific painting, a powerful tool for revealing karyotype evolution, could provide further insights into the structure of the YD/Neo-Y (Han et al., 2015).

In recent years, technical advances in DNA synthesis have allowed massively parallel de novo production of thousands of independent oligonucleotides (oligos) reducing their cost (Han Y. et al., 2015). The OligoMiner pipeline allows high-throughput design of specific single-copy oligo probes, successfully labeling chromosomal regions, arms, or entire chromosomes in humans and *Drosophila* (Beliveau et al., 2018; Rosin et al., 2018). However, the minimal nucleotide divergence between the neo-sex chromosomes (Zhou et al., 2012) and the highly repetitive nature of the YD/Neo-Y chromosome limits the density of single-copy oligo probes achievable for fluorescence *in situ* hybridization (FISH).

Here, we exploited the most recent genome assembly of *D. miranda* (Mahajan et al., 2018) to design single-copy and repetitive-oligos libraries for painting the XL/XR, Neo-X, YD/Neo-Y sex chromosomes, and the autosome 4 (Muller B) (Fig 1C; S1 Fig). These libraries were achieved by integrating OligoMiner with the approach suggested on the OligoY pipeline (Almeida et al., 2025). Oligo specificity was tested via fluorescence *in situ* hybridization (FISH) on larval neuroblast chromosome spreads, and we analyzed oligo density (*hits*/kb) and spacing to predict FISH labeling success. By physically mapping the three largest Y linked scaffolds (Y1, Y2, and Y3) of *D.miranda* and using these libraries in the sister specie *D. pseudoobscura*, we shed light on unprecedented events that gave rise to the *D. miranda*’s Y chromosome: chromosomal translocation, centromere loss, inversion and DNA amplification involving ancestral YD and the Muller C element. Our oligo libraries are the first designed for *D. miranda* and the only ones available in the *Drosophila* genus besides *D. melanogaster*. As chromosome identification is crucial for cytogenetic research, these libraries provide a valuable resource for studying sex chromosome behavior across different cell types.

## Results

### Single-copy oligos effectively label autosomes and X’s chromosomes of different ages

The specific targeting of the Neo-X chromosome with only 1.5 million years (Mahajan et al., 2018) indicates that, despite the high similarity with its Neo-Y homolog (Zhou et al., 2012), there are sufficient nucleotide differences to design specific oligos. Therefore, we have fully painted euchromatic regions of the Neo-X, as well as the 4th and XL/XR chromosomes (ancestral X, 60 Ma; XR-D, 15 Ma) using single-copy oligos (Fig 2A, 2B, 3C, S2 Fig).

**Fig 2.**
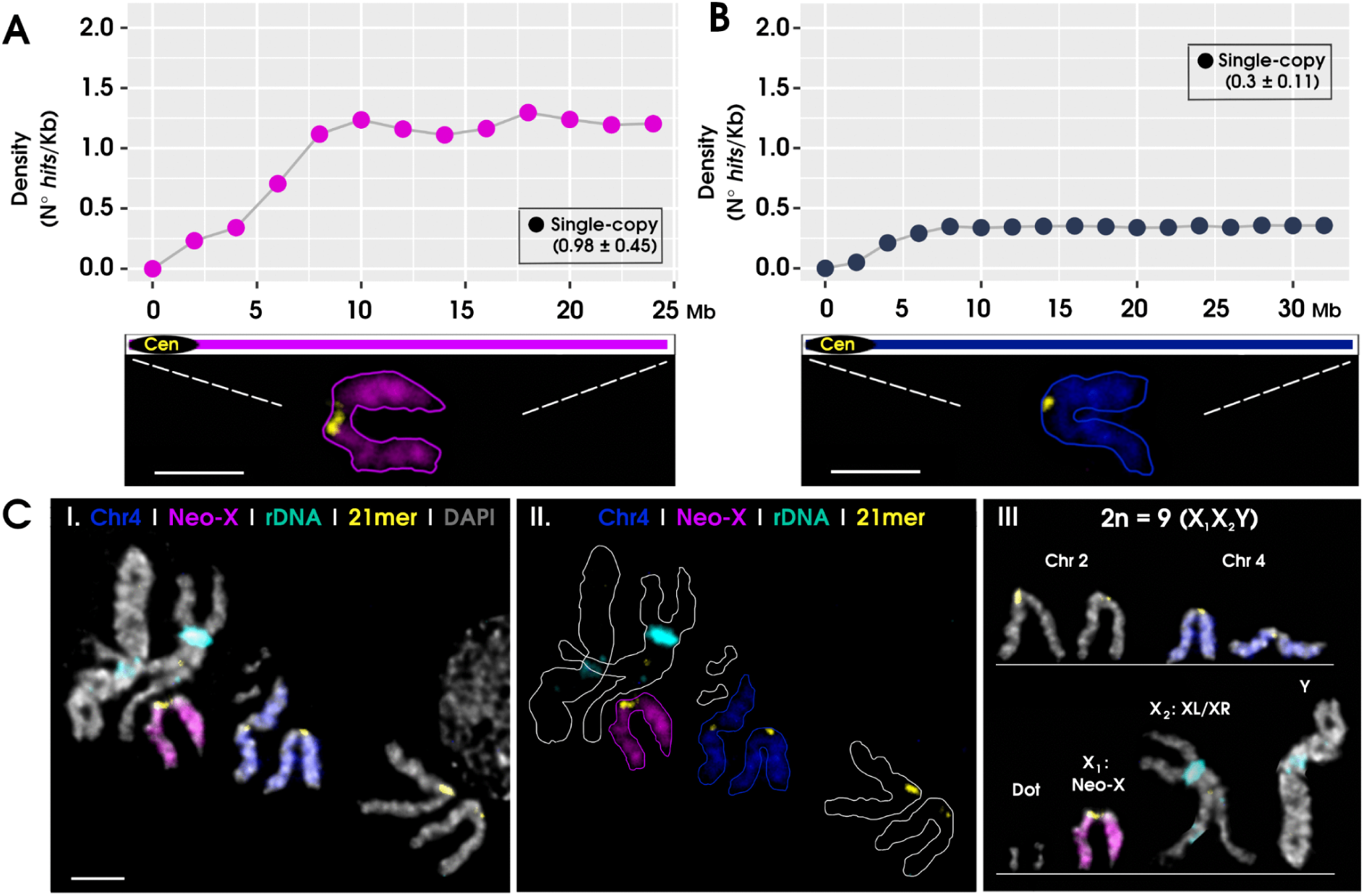
Painting of Neo-X and Chromosome 4 in *D. miranda*. **A)** and **B)** Show the oligo density (no. *hits*/kb) for single-copy oligos on Neo-X and Chromosome 4, respectively. The x-axis represents the DNA sequence position on each chromosome. Oligo density was computed using a 2Mb sliding window (mean ± SD). **C)** Male metaphase (2n=9) labeled with single-copy oligos targeting Neo-X (Cy5: magenta) and Chromosome 4 (Cy5: blue), rDNA (anti-DIG-rhodamine: cyan), and centromeric 21mer (Avidin-FITC: yellow) probes. **C.II)** Shows the labeling on chromosomes with only their contours outlined. **C.III** represents the karyotype for male metaphase. X_1_: Neo-X, X_2_: XL/XR. Cen: centromere. Bar scale, 2.5 μm.

Repetitive DNA regions, like centromeric, pericentromeric, or ribosomal DNA, often lack enough single-copy oligos, resulting in weak or absent labeling, as seen near centromeric regions (Fig 2A, 2B, 3A). To address this, we first determined the oligo density threshold needed for effective labeling across all regions studied, both here and in the following sections, optimizing the process and conserving resources.

Studies with single-copy oligos have shown that required density varies by target size: smaller regions need higher densities, while larger regions can be labeled with lower densities. Typically, kilobase, megabase, and whole chromosome regions require minimum densities of 2-3.4 *hits*/kb, 1.5 *hits*/kb, and 0.25-1 *hits*/kb, respectively (Yamada et al., 2011; Beliveau et al., 2015; Albert et al., 2019; Rosin et al., 2018). Our findings align with these results. We achieved effective labeling at 1 hit/kb for whole chromosomes or 25 Mb regions, such as Neo-X (Fig 2A), XR-D, and XL-2 (Fig 3B, C). Even lower densities of 0.32 *hits*/kb were sufficient to label chromosome 4 (32 Mb), which was slightly higher density than the 0.25 *hits*/kb used for 160-300 Mb maize chromosomes (Albert et al., 2019).

**Fig 3.**
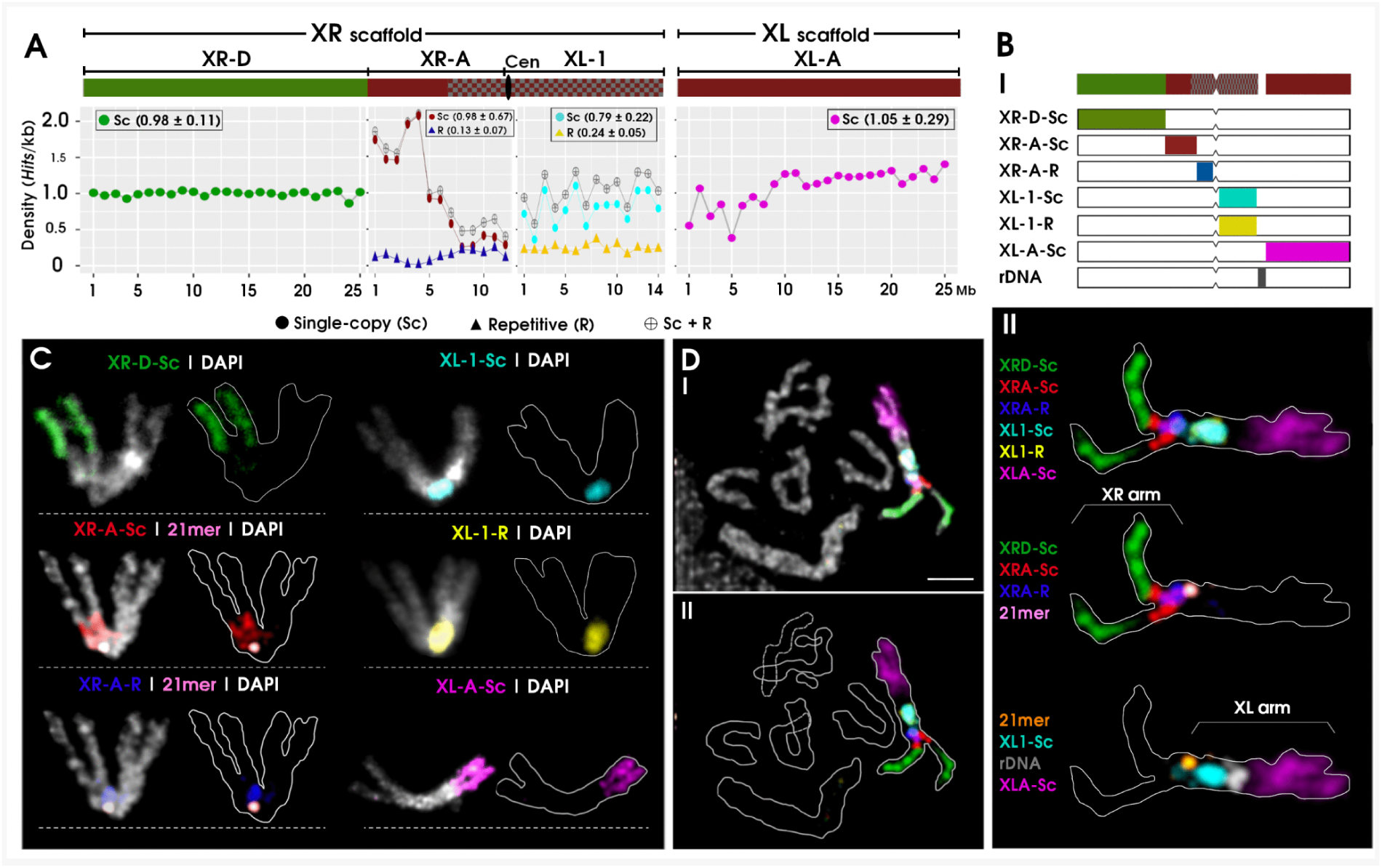
Painting of the XL/XR chromosome of *D. miranda*. **A)** Density (no. *hits*/kb) of single-copy (Sc), repetitive (R) and combine (Sc+R) oligos along the three regions of XR scaffold (XR-D, XR-A, XL-1) and the XL scaffold (XL-A). The x-axis shows chromosome position (Mb). Density of oligos was computed using a window of 1Mb (mean ± SD). **B)** Mapping of the oligopaints libraries on the XL/XR chromosome. **B.I** Schematic representation of the oligopaints’s labeling and rDNA probe. **B.II** Multiple oligo-FISH with our six oligopaints libraries, rDNA and centromeric (21mer) probes painting the complete chromosome. XR-D-Sc (Cy5: green), XR-A-Sc (atto488: red), XR-A-R (atto488: blue), XL-1-Sc (Cy5: cyan), XL-1-R (Cy5: yellow), XL-A-Sc (atto488: magenta), rDNA (antiDIG-rhodamine: gray) and centromeric (Avidin-FITC: pink or orange). **C)** Individual mapping of every oligo library along the XL/XR chromosome. **D)** Male metaphase labeled with the six oligopaints libraries showing no off-target hybridization. **D.II** Oligopaints label on XL/XR chromosomes with only their contours outlined. Bar scale, 2.5 μm.

### Assessing the XL/XR assembly using oligo-FISH

The most recent genome assembly for *D. miranda* failed to combine the XL and XR chromosomes into a single continuous scaffold, leaving them as two separate scaffolds corresponding to XL and XR chromosomes (Mahajan et al., 2018; Fig 3A). Our oligo-FISH and rDNA probe analyses revealed that rDNA is located between these scaffolds (Fig 3B). Combined with the similar rDNA localization pattern in *D. pseudoobscura*, which shares the same chromosome (Larracuente et al., 2010), this suggests that the tandem repetitive rDNA structure caused the assembly failure (McKee et al., 1992).

In short, the XL/XR chromosome was fully painted using our single-copy and repetitive oligos libraries, along with the rDNA probe (Fig 3B). While most regions along the XL/XR sequence show single-copy oligo densities around 1 hit/kb (Table A in S1 Text), the distribution is uneven (Fig 3A). For example, the pericentromeric region of XR-A (7-12 Mb) has densities from 0.25 to 0.5 *hits*/kb. In these cases, assessing inter-oligo spacing can help predict labeling effectiveness. Within this region, the median spacing is about 1000 nucleotides (1 hit/kb), though variability is high (S3 Fig). Similar patterns occur in the centromeric areas of the neo-X and chromosome 4 (S4 Fig), leading to weak or absent labeling. Interestingly, when spacing distribution is narrow and centered around the median, labeling is still possible even with oligos spaced beyond 1000 nucleotides, as seen in chromosome 4 (S4 Fig).

To tackle weak or absent labeling due to low-density single-copy oligos, we adapted our Oligo Y pipeline (Almeida et al., 2025). Previously, we enhanced the OligoMiner workflow with specific repetitive oligos, boosting oligo density for the *D. melanogaster* Y chromosome by over 100-fold (Almeida et al., 2025). Here, we applied this approach to design exclusive repetitive oligos for the XL/XR and Y chromosomes of *D. miranda* (See Generation of repetitive oligopaints in Materials and Methods).

Repetitive oligos are present in regions with few single-copy oligos, as shown by labeling the pericentromeric region of XR-A, comparing FISH results from the XRA-Sc and XRA-R libraries (Fig 3C, S5 Fig). In the XL-1 region, despite differences in single-copy (0.78) and repetitive (0.23) oligo densities (Table A in S1 Text), both libraries produce similar labeling patterns (Fig 3C, XL1-Sc and XL1-R). This can be explained by the repetitive library’s narrow inter-oligo spacing distribution centered around the median (S3 Fig, D-II), suggesting that a 1 hit/kb density cut-off is not necessary for effective labeling.

### The Y chromosome is fully labeled using single-copy and repetitive oligo probes

The *D. miranda* Y chromosome consists of two regions: YD (15 Ma) and Neo-Y (1.5 Ma). The latest genome assembly identified three main Y chromosome scaffolds: Y1, Y2, and Y3. Scaffolds Y1 and Y2 correspond to the Neo-Y, showing similarity to the Muller C element, while Y3 aligns with the Y chromosome of *D. pseudoobscura*, representing the YD region (Mahajan et al., 2018).

Our approach based on OligoY successfully identified specific repetitive oligos, completing the labeling of the entire *D. miranda* Y chromosome (Fig 4) and demonstrating the method’s broad applicability across different evolutionary stages of Y chromosomes. Although the oligo densities for both single-copy and repetitive libraries were below 1 hit/kb (ranging from 0.25 to 0.59 *hits*/kb), the Y1 and Y2 scaffolds exhibited uniform labeling across their entire lengths (Table A in S1 Text, Fig 4A). The inter-oligo spacing was centered around the median, consistent with the pattern observed in other fully labeled chromosomes (S6 Fig). Regarding centromere positioning, the Y1 scaffold is located on the left arm, extending toward the proximal region near the centromere on the right arm, whereas the Y2 scaffold is exclusively localized on the right arm (Fig 4B, D).

**Fig 4.**
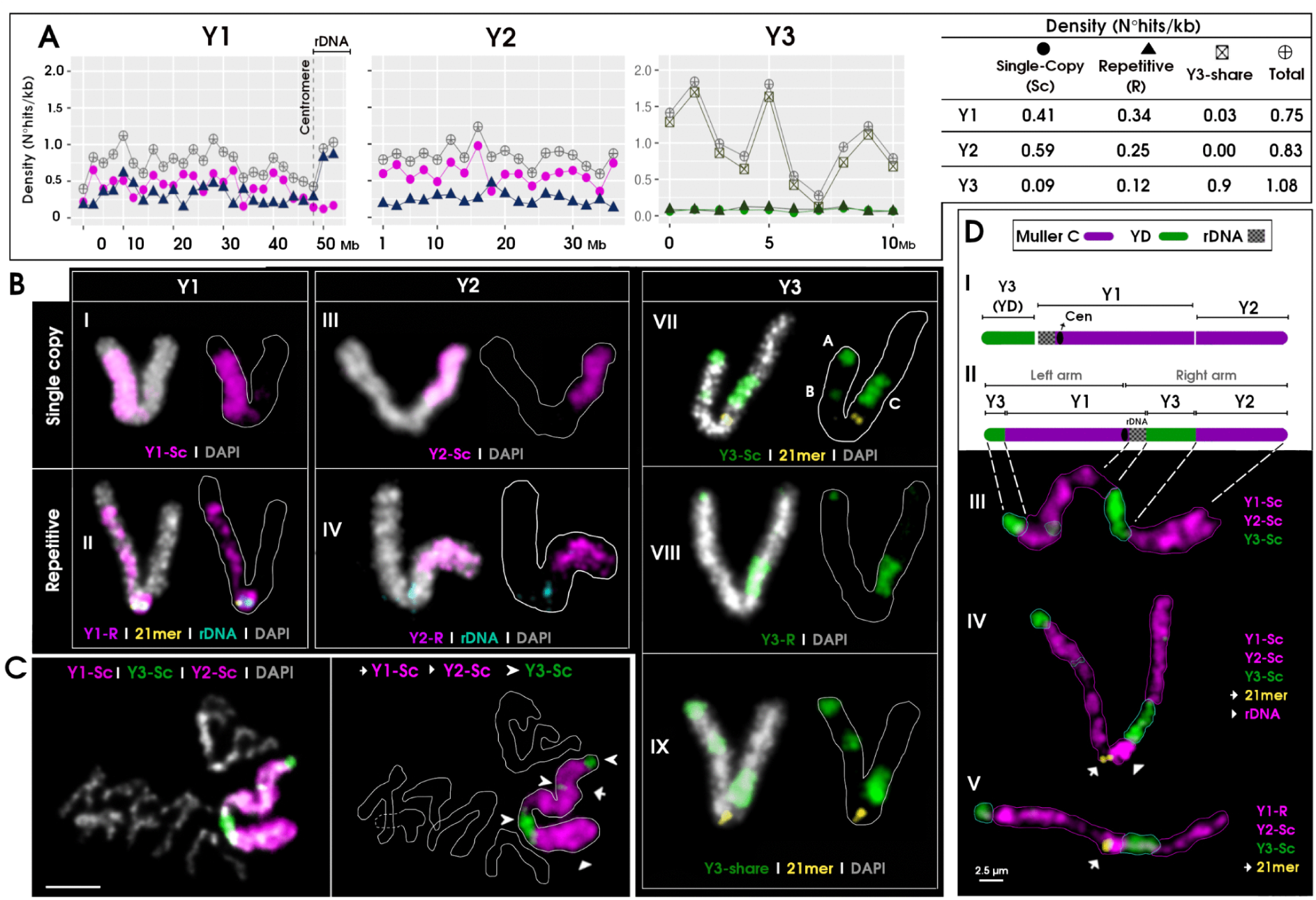
Mapping of three major Y-linked scaffolds (Y1, Y2 and Y3) on the Y chromosome of *D. miranda*. **A)** Density (no *hits*/kb) of single-copy (Sc), repetitive (R) and total (Sc+R) oligos along the Y1, Y2 and Y3. For Y3 scaffold, density of Y3-share and combined (Sc+R+Y3-share) is also shown. Density of oligos was computed using a window of 2Mb for Y1 and Y2, and 1Mb for Y3. Values of density of oligos for scaffold are presented in a table. **B)** Oligo-FISH using single-copy oligos for Y1 (Cy5: magenta), Y2 (Cy5: magenta), Y3 (Cy5: green), and repetitive oligos for Y1 (Cy5: magenta), Y2 (atto488: magenta), Y3 (atto488: green), Y3-share (atto488: green), and rDNA (antiDIG-rhodamine: cyan) and 21mer (Avidin-FITC: yellow) probes. B.V. The letters: A, B and C represent the three labels of Y3-Sc oligos in the Y chromosome. **C)** Male metaphase with single-copy oligos for Y1, Y2 and Y3 showing no off-target hybridization of Y-linked oligos. **D)** Structure of Y chromosome. **D.I** represents the Y according to Mahajan S. et al., (2018). **D.II** Schematic representation of the Y according to our FISH results. Note that the Neo-Y (Muller C: Y1 and Y2) region is interspersed by Y3 (YD). **D.III** to V show that the Y chromosome is fully labeled by using our oligo libraries.

The rDNA probe targets a region adjacent to the centromere on the Y chromosome, situated at the end of the Y1 scaffold and exclusively labeled by the repetitive Y1 oligo library (Fig 4A, 4B-II). Notably, Y1-R labels rDNA on the Y chromosome but not on XL/XR (S7 Fig), confirming its specificity for Y-linked rDNA sequences.

In addition, FISH targeting the Y3 scaffold produced intense fluorescence despite low oligo densities (0.09–0.12 *hits*/kb; Table A in S1 Text), likely due to underestimation, as repetitive oligos may hybridize to multiple loci along the repetitive scaffold. A similar pattern was observed on the *D. melanogaster* Y chromosome (∼60 Ma), which showed strong labeling with only 0.13 *hits*/kb (Almeida et al., 2025). Indeed, by increasing search sensitivity and adjusting alignment criteria in the Bowtie2 analysis (see ‘Enhancing Oligo Density Estimation’ in Materials and Methods), the estimated Y3 oligo density increased by up to 3.1-fold, reaching a level sufficient for FISH detection. Specifically, the single-copy and repetitive Y3 libraries increased to 0.26 and 0.39 *hits*/kb, respectively, with sufficient levels for fluorescence detection and still comparable to those of other chromosomes. In contrast, applying the same enhanced search sensitivity did not alter the oligo densities for single-copy libraries on the autosomes, neo-X, or XL/XR (S2 Table).

### Assessing *D. miranda* Y chromosome scaffolds assembly

Unexpectedly, oligo libraries for scaffold Y3 label three distinct regions (A, B, and C) on the Y chromosome of *D. miranda* (Fig 4D, V), despite being designed for a single continuous scaffold. Chromosomal measurements indicate that label B is located around 17.15 ± 2.1 Mb on scaffold Y1 (S8 Fig, A-III). Further alignment using an enhancing criteria (refer to Enhancing Oligo Density Estimation in Materials and Methods) revealed that some Y3 oligos also aligned around this position (S8 Fig). These oligos align within the FBgn0082139 gene, which is repeatedly found at 17 Mb on scaffold Y1 (Bachtrog et al., 2019). FBgn0082139, orthologous to *D. melanogaster’s fest*, is annotated on both Y3 and Y1, possibly explaining the cross-alignment of certain Y3 oligos to Y1 (Bachtrog et al., 2019).

The remaining two labels, A and C (Fig 4D-VI), are unlikely to be off-target signals, as they do not overlap with oligo labeling for scaffolds Y1 or Y2. How could *D. miranda’s* Y chromosome have two distinct, yet separated, regions homologous to YD? A plausible hypothesis is that a large-scale inversion occurred after chromosomal fusion, disrupting the YD chromosome (Fig 5A). To test this, we employed two approaches: (1) cross-species oligo-FISH using *D. miranda* oligos to label *D. pseudoobscura* chromosomes, in which Muller C (chromosome 3) remains autosomal rather than forming a neo-sex chromosome; and (2) large-scale sequence alignment between Muller C of *D. pseudoobscura* and the Y1 and Y2 scaffolds to identify potential structural rearrangements.

**Fig 5.**
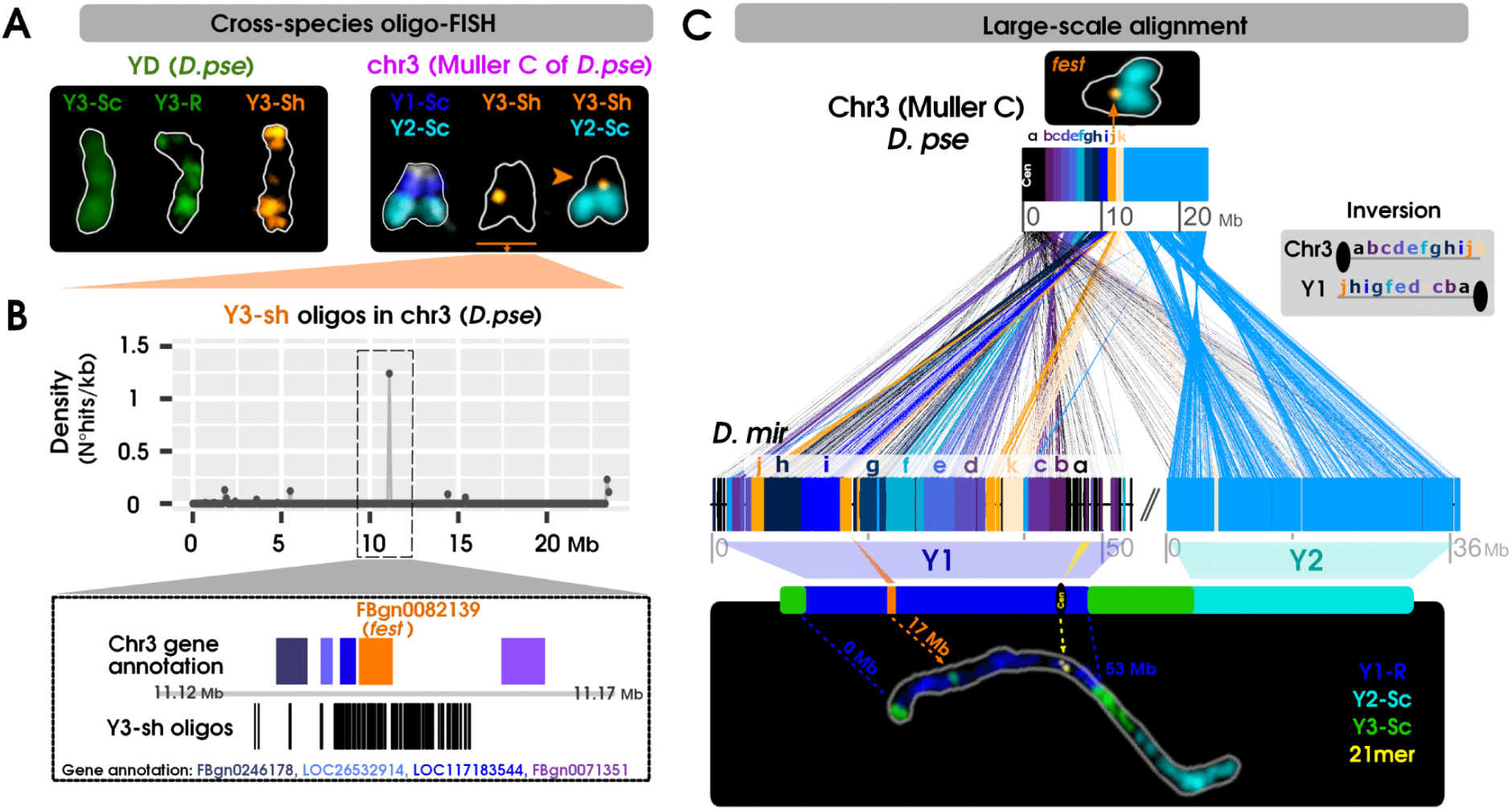
Investigating large-scale rearrangements shaping the Y chromosome in *D. miranda*. **A)** Cross-species oligo-FISH in *D. pseudoobscura* using *D. miranda* oligo libraries. **B)** Interstitial labeling on Muller C (chr3) by Y3-sh library; oligo density peaks near 11 Mb, aligning with the FBgn0082139 gene, orthologous to *fest* in *D. melanogaster*. **C)** Large-scale alignment between *D. pseudoobscura* Muller C and *D. miranda* scaffolds Y1 and Y2. Cross-species FISH results with Y3-sh mark *fest* as the approximate breakpoint between Y1 and Y2 in *D. miranda* (orange mark). The region from the centromere to *fest* was divided into 1 Mb segments (coloured a–k), revealing a large-scale inversion on Y1 through inverted positions of these segments in *D. miranda*.

Cross-species FISH experiments revealed that the Y1 and Y2 oligos specifically labeled continuous regions on Muller C in *D. pseudoobscura*: Y1 oligos marked the region from the middle to the centromere, and Y2 oligos extended toward the telomere (Fig 5A, S9 Fig), supporting the hypothesis that Y1 and Y2 correspond to the neo-Y (Mahajan et al., 2018). Additionally, the labeling of the *D. pseudoobscura* Y chromosome by both single-copy and repetitive Y3 oligos suggests a similarity between scaffold Y3 and the ancestral YD chromosome (Fig 5A, S9 Fig, B and C). In contrast, the library Y3-sh labeled autosomal telomeres and an interstitial signal on chr3, positioned between Y1 and Y2 boundaries, corresponding to FBgn0082139, an ortholog of the *fest* gene in *D. melanogaster* (Fig 5B, S9 Fig, D–F). This approximately marks the breakpoint on *D. pseudoobscura* Muller C that gave rise to the Y1 and Y2 scaffolds in *D. miranda*.

Large-scale alignment revealed that the Y1 scaffold sequence is inverted on the *D. miranda* Y chromosome (Fig 5C). However, two internal regions within Y1, segments *h* and *i*, remain non-inverted. Additionally, segment *k*, which corresponds to the breakpoint boundaries on chromosome 3, is now located internally within Y1. These observations suggest that multiple inversions likely contributed to the structural reorganization of the Y chromosome during its evolution.

We then investigated what label A represents. The *fest* gene, located on Y1 and Y3, has amplified to groups of 1, 2, and 24 copies annotated at 0.6 Mb, 6.7 Mb, and 17.1 Mb on Y1 (Bachtrog et al., 2019). Given that the chromosomal breakpoint occurred near the *fest* gene, the Y1 inversion was initially thought to explain the signal at 0.6 Mb near label A, which was presumed to mark the ancestral *fest* locus. To test this, we designed oligo probes targeting the *fest* gene of *D. pseudoobscura*. FISH experiments showed that the probes hybridized exclusively to label B within Y1, demonstrating that label A does not correspond to the *fest* gene (S10 Fig). Another possibility is that label A corresponds to the Y6 scaffold, which showed sufficient oligo density for FISH detection using Y3-derived libraries among the other Y-linked shorter scaffolds (Table C in S1 Text). The Y6 harbors 24 copies of the gene *FBgn0245922* (Bachtrog et al., 2019). This gene, an ortholog of *pain* in *D. melanogaster*, is also amplified on scaffolds Y1 (47 copies) and Y3 (90 copies). Notably, the Y6 scaffold contains telomeric sequences, supporting its likely localization at the distal end of the left chromosomal arm. To test this hypothesis, we designed oligo probes targeting the *pain* gene. FISH confirmed that label A represents the *pain* gene (S10 Fig). Since, the *pain* gene is originally located on the autosomal Muller C of *D.pse,* this indicates that label A corresponds to Muller C rather than YD.

### *D. miranda* Y Chromosome: translocation, centromere loss and rearrangements

Our findings support a model for the evolutionary events that shaped the Y chromosome of *D. miranda* (Fig 6). We suggest that the process began with a translocation of a segment from the ancestral YD chromosome to the telomeric region near the centromere of the short arm of Muller C. It remains unclear whether this translocated segment originated from the long or short arm of YD. This event likely gave rise to a metacentric chromosome, with interstitial telomeric sequences positioned centrally on the Y chromosome. Supporting evidence includes the presence of telomeric oligos from the Y3-sh library (Fig 4B-VII) and previous telomeric probe labeling (Steinemann et al., 1984), both marking the central region of the *D. miranda* Y chromosome. As a consequence of this translocation, two Y chromosomes were initially formed; however, the unfused YD chromosome, still carrying a centromere, was likely lost to ensure meiotic stability.

**Fig 6.**
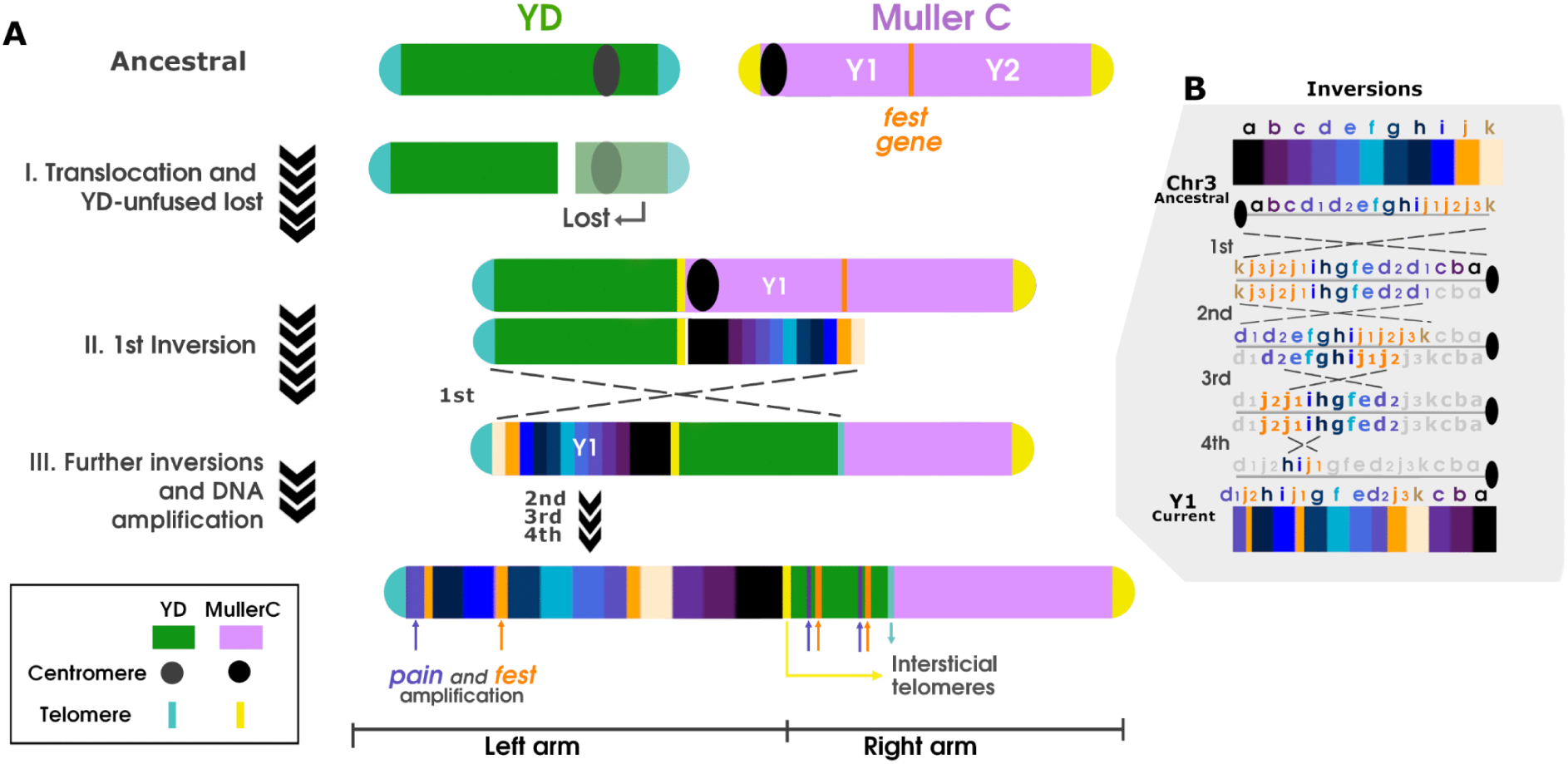
Stepwise Evolution of the *Drosophila miranda* Y Chromosome. **A)** The current architecture of the *D. miranda* Y chromosome can be explained by three major evolutionary steps. **Step I:** A segment from the long arm of the YD chromosome translocated to a telomeric region near the centromere of Muller C, while the remaining YD portion was subsequently lost. **Step II:** A large-scale pericentric inversion breaking around the *fest* gene in Muller C rearranged the YD translocated segment and Y1. **Step III:** Additional inversions and extensive DNA amplification further modified the structure, giving rise to the current organization of the Y chromosome. FISH results indicate that the *fest* gene expanded from its original position at the end of the left arm to the Y1 and Y3 scaffolds. **B)** Four inversions within the Y1 scaffold recapitulate its current organization and explain the internal positioning of genomic regions that were originally distal.

Large-scale alignments indicate that four sequential inversions shaped the present structure of the *D. miranda* Y chromosome (Fig 6B). Following the translocation, a pericentric inversion occurred involving regions of Y1 and YD, resulting in an interspersion of Y1 and Y2 sequences with those from YD. The inversion breakpoint appears to lie near the *fest* gene, which was repositioned toward the distal end of the left arm (Fig 6B, region j in orange). In YD, this inversion, encompassing most of the entire translocated segment, may have repositioned part of the YD telomere to the middle of the chromosome. As a result, the interstitial telomeric sequences likely originated from both YD and Muller C, while the centromere of the current Y chromosome was most likely inherited from Muller C (Fig 6A). The second inversion, spanning the region between *k-d1*, accounts for the internal positioning of *k* within Y1, as revealed by the large-scale alignment (Fig 5C). This inversion also accounts for the present position of the *pain* gene. This gene originally located in 5,6 Mb in Muller C of *D. pse* represented by region *d* is partially repositioned at the beginning of the Y1 scaffold (See the purple d region at the start of Y1 in Fig 5C). The third inversion, occurring between d2-j2, split the *d* and *j* regions, placing *j3* adjacent to *k*. The fourth inversion, involving the *j1-h* region, explains the conserved orientation of this segment between Y1 and ancestral chromosome 3 (Muller C) of *D. pseudoobscura*. Notably, this inversion clarifies why *fest* copies are clustered near 17 Mb rather than closer to the Y1 terminus, as would have been expected after the pericentric inversion. Furthermore, the *fest* copies at *j2*, along with its annotations around ∼6.7 Mb (Batchrog et al., 2019) indicated that this gene amplification began prior to the fourth inversion, which subsequently split the *fest* copies. Thus, the Y chromosome experienced extensive DNA and gene amplification both during and after the inversions that shaped its current architecture.

An alternative scenario in which the YD translocation involved the long arm of Muller C is less likely. Such an event would have produced an acrocentric Y chromosome, and even if followed by an inversion to explain the discontinuous arrangement of Y1 and Y2, it would still not account for the current structure of the *D. miranda* Y (S11 Fig).

## Discussion

This study demonstrates the successful design of oligo libraries for *Drosophila miranda*, particularly targeting its sex chromosomes with evolutionary ages ranging from 1.5 to 60 million years ago (Mahajan et al., 2018). Our results highlight the versatility and applicability of these libraries, offering valuable tools for various studies. Specifically, we were able to design effective oligos for both single-copy and repetitive regions, which is especially important for investigating several cellular processes. For instance, sex chromosomes often exhibit distinct structural and functional roles during meiosis (Vibranovski 2014), and our oligos will enable precise labeling and analysis in these contexts.

Through the design and testing of oligos, we uncovered several key insights. First, the density of oligos is a critical parameter for successful chromosome painting. While previous studies with *D. melanogaster* used densities of 1 oligo per kilobase (Rosin et al., 2018; Mahadevaraju et al., 2021), we found that densities as low as 0.5 oligos per kilobase are sufficient, provided that the inter-oligo spacing does not exceed 2,500 nucleotides. This optimization allows for cost-effective labeling while maintaining robust fluorescence signals. Second, we demonstrated the cross-species reproducibility of oligos, with those designed for *D. miranda* successfully labeling homologous regions in *D. pseudoobscura*. This finding suggests that our method can predict the applicability of oligo libraries in related species, facilitating broader evolutionary and comparative genomic studies. Third, the method’s effectiveness in repetitive regions, such as pericentromeric and centromeric DNA, expands its utility. Repetitive oligos, requiring shorter libraries, achieve fluorescence levels comparable to single-copy oligos, as observed in the XL-1 region (Fig 3C). These findings demonstrate that repetitive oligos represent not only a viable alternative but also a resource-efficient approach for investigating chromosome structure and behavior

By designing these oligos we gained unprecedented resolution of the *Drosophila miranda* Y chromosome assembly, underscoring the importance of oligopaint for uncovering true scaffold configurations. Using these tools, we mapped the three major Y chromosome scaffolds, Y1, Y2, and Y3, originally identified through genome sequencing (Mahajan et al., 2018). Our analyses revealed that Y1 and Y2 are located on opposite chromosomal arms, with Y3 occupying the intervening region. Whereas a simple fusion of Muller C and YD would predict Y1 and Y2 to be contiguous, the placement of Y3 (derived from YD) points to a more complex evolutionary history. This configuration also explains why Y1 and Y2 failed to assemble into a single scaffold during previous genome assemblies.

We propose that a pericentric inversion following the translocation of YD to Muller C might reposition the entire YD to the central region of the Y chromosome. Alignments between the neo-Y (Y1 and Y2) of *D. miranda* and Muller C of *D. pseudoobscura* provide evidence of inversions in Y1, consistent with patterns of inversions concentrated in Muller C (chr3) of *D. pseudoobscura* (Dobzhansky & Sturtevant, 1938; Wallace et al., 2011). Similar processes, including fusions followed by pericentric inversions, have shaped neo-sex chromosomes in species such as *D. albomicans* and *D. busckii* (Wang et al., 2022; Krivshenko, 1955). Given the composition of the *D. miranda* Y chromosome, it is possible that transposable elements (TEs) and short repeat sequences contributed to these genomic rearrangements, as they have been identified as driving forces in closely related species (Cáceres et al., 1999; Richards et al., 2005).

The Y centromere of *D. miranda* likely originated from Muller C but was invaded by a 99mer satellite within the last 1–2 million years (Bracewell et al., 2019). While rDNA is absent in Muller C and only intergenic spacer regions (IGS) are present in YD of *D. pseudoobscura* (Larracuente et al., 2010), the *D. miranda* Y chromosome contains rDNA near the centromere. This rDNA may have been acquired from the XL/XR chromosome through translocation with posterior amplification driven by transposable elements (TEs). Supporting this, copies of R2 retrotransposon, essential for expansion of rDNA in the *Drosophila* male germline (Nelson et al., 2023), are found in the Y chromosome of *D. miranda* by blast alignment. Furthermore, the R1 retrotransposon, found in the 28S region of rDNA across the *Drosophila* genus, is also present in centromeric heterochromatin (Eickbush et al., 1997). The rDNA translocation likely created double-strand breaks near the centromere, facilitating the formation of the 99mer satellite, as CENP-A, a centromere-specific histone H3 variant, is known to be recruited to such breaks (Zeitlin et al., 2009), enabling the emergence of a neo-centromere (Bracewell et al., 2019).

A key question for future research is identifying which region of the YD chromosome was translocated to Muller C. While we hypothesize that the long arm was involved, the possibility of a short-arm translocation cannot be ruled out. Unfortunately, the highly repetitive nature of the *D. pseudoobscura* YD chromosome has thus far precluded the development of a chromosome-level assembly, limiting our ability to design informative oligo probes.

## Materials and Methods

### Fly strain

We used the inbred MSH22 and 14011-0121.94 (Drosophila Species Stock Center code) strains for *D. miranda* and *D. pseudoobscura* respectively. Both species were maintained in a chamber at a constant temperature of 18°C, with consistent humidity and a photoperiod of 12h:12h light/dark. The MSH22 *D. miranda* was kindly donated by Kevin Wei from the Bachtrog Lab at UC Berkeley.

### Identification of Y-linked scaffolds

The reference genome of *D. miranda* (Genbank GCA_003369915.2), which consists of 104 scaffolds, was used to identify Y-linked scaffolds using the combined results of two approaches: Y-Genome Scan (YGS) based on k-mer (Carvalho et al., 2013) and the suggested classification done during genome assembly based on coverage male/female reads (Mahajan et al 2018). Briefly, female and male short reads (Genbank SRR789669, SRR364796) were used to run YGS employing *k*-mers 15 and 18. 15-mers are typically used to filter errors in short reads from Drosophila sequencing, and 18-mers can enhance the resolution for classifying contigs. Then, scaffolds with a proportion of unmatched female single-copy k-mers > 0.8 were considered Y-linked (Carvalho et al., 2013). Finally, we intersected the three lists of classifications (15-mers, 18-mers and coverage male/female reads) to identify Y-linked scaffolds. Using this method, 41 scaffolds were identified as Y-linked including the largest Y1, Y2 and Y3 scaffolds. Moreover, the chromosomes chr2 (Muller-E), chr4 (Muller-B), chr5 (Muller-F), neo-X (Muller-C), XL (Muller-A, refers as XL-A), XR (Muller-AD) were undoubtedly classified as X or autosome (X/A) as was expected (S12 Fig).

### XR scaffold structure and scaffolds definition

Given that the XR scaffold (Mahajan et al 2018) encompasses Muller A and D elements and extends into a region on the left arm of the XL/XR chromosome, its structure has been segmented based on Muller element similarity and the centromere’s position. For that, Muller element’s similarity to *D. melanogaster* (Release 6) and the 99-mer (5’AAAAAATCACTTTGCACCAAACTCATTTTATGCCTAAATCGACCACATATTTAGCTTATTTGTACCCAAAATCGACCAAAAACCGACACAATTTTAAC3’) and 21-mer (5’AAAACTTATCTCCGCTGGCGG3’) tandem centromeric repeats position (Mahajan et al 2018) were analyzed using Nucmer (from MUMmer software, Kurtz et al., 2004) and BLAST (Altschul et al., 1990) respectively. Thus, XR scaffold was divided into three regions called XR-D (region with similarity to Muller D in the XR arm), XR-A (region with similarity to Muller A in XR arm) and XL-1 (region located in XL arm) (Fig 3A). Consequently, our final genome used to design oligopaints was composed of 106 scaffolds.

### Generation of single copy oligopaints

Single-copy oligos were designed using OligoMiner (Beliveau et al., 2018) for chr4, neo-X, XR-D and XL-2 (S1 Fig). Briefly, the sequences were masked by RepeatMasker (Smit et al., 2015) and used for probe discovery. BlockParse script was run considering the stringent set designed to maximize probe-binding affinity (40–46 nt, 47-52 °C hybridization) and overlap (-O) to retrieve all possible oligos. Bowtie2 was run with “--very-sensitive-local –k 2 -t” using Bowtie2 alignment indices builded with the genome assembly (106 scaffolds) without repeat masking. The outputClean script was run with default values to retrieve unique oligos (-u). Postprocessing steps included K-mer, structure check and density reduction filters to increase specificity, predict secondary structure and select oligos spaced by a desired nucleotide distance respectively. Thus, a 18-mer count was generated for the genome assembly (106 scaffolds) without repeat masking using “-C -m 18 -s 10G -t 3” using Jellyfish (Marçais et al., 2011). The kmerFilter script was run considering 18-mer and “-k/–kmerThreshold” set to 5. The structureCheck script was used with the default setting “-T 37 -t 0.1”. The DensityReduction script was run to retrieve 1 oligo/kb testing several nucleotide distances between oligos. Density of 1 oligo per kilobase (kb) is commonly employed for chromosome painting in *D. melanogaster* (Rosin et al., 2018). However, in the case of chr4, we opted for a lower density of 0.34 oligos/kb to investigate the efficacy of a reduced density in chromosome painting. A lower density value represents a cost-saving measure for future experiments. The resulting BED file was used for statistical analysis of oligos along the target scaffold using R software and probe construction.

### Generation of repetitive oligopaints

Repetitive-oligos were designed using a modification of the conventional pipeline of OligoMiner (Beliveau et al., 2018) suggested in OligoY (Almeida et al., 2025) for XR-A, XL-1, Y1, Y2 and Y3 (S1 Fig). XR-A and XL-1 represent the scaffolds located on both sides of the centromere in XL/XR, and Y1, Y2, Y3 are YD/neo-Y linked scaffolds. The sequences without repeat masking were used for probe discovery. BlockParse script was run considering the stringent set (40–46 nt, 47-52 °C hybridization) and overlap (-O) to retrieve all possible oligos. Bowtie2 was run two times: (1) using Bowtie2 alignment index builded with the complete genome assembly (106 scaffolds) and (2) using Bowtie2 alignment index builded with the genome assembly without the sequence target (105 scaffolds). Since the genome does not contain the target sequence, oligos that fail to align can be regarded as oligos for the target sequence. Bowtie2 was run with “--very-sensitive-local –k 2 -t”. Next, the outputClean script was run with default values for the two previous results. For (1) was used “-u” to retrieve unique (single-copy) oligos and for (2) was run “-0” to select oligos that failed to align. These oligos represent the complete list of oligos comprising both single-copy and repetitive-oligos. Thus, repetitive-oligos were obtained by subtracting single-copy oligos (result 1) from the complete list of oligos (result 2). Postprocessing steps included K-mer, structure check, female short reads and, finally, density reduction or a combined density reduction. Female short reads filter is added only for Y linked scaffold targets. Density reduction step is used when the user is interested in only repetitive-oligos whereas combined density reduction step is chosen for obtaining single-copy and repetitive-oligos. The CombinedDensityReduction script uses both lists of oligos to select single-copy oligos spaced by a desired distance (nt) and incorporates repetitive-oligos in positions where single-copy oligos are absent, while also considering a specific spacing with the preceding oligo. Postprocessing steps were independently run for the single-copy and repetitive-oligos as follows: For single-copy oligos, a 18-mer count was generated for the genome assembly (106 scaffolds) without repeat masking using “-C -m 18 -s 10G -t 3” in Jellyfish. The kmerFilter script was run considering 18-mer and “-k/–kmerThreshold” set to 5. For repetitive-oligos, a 18-mer count was generated for the genome assembly without the target sequence (105 scaffolds) and kmerFilter script was run as described before. The structureCheck script was used with the default setting “-T 37 -t 0.1” for single-copy and repetitive-oligos. Only for Y-linked oligos, female short reads (forward and reverse) (Genbank SRR789669) were used to build Bowtie2 alignment indices, and Bowtie2 was run to align single-copy and repetitive-oligos with “--very-sensitive-local –k 2 -t”. The outputClean script was run with “-0” to select oligos that failed to align. Since the Y chromosome is absent in female short reads, this additional filter increases further the Y linked specificity of our oligos. Finally, a CombinedDensityReduction script was run testing different nucleotide distances between oligos to reach 1 oligo/kb for XR-A and XL-1. This value of density could not be reached for Y-linked scaffolds, thus the CombinedDensityReduction script was run with “-nU 0 -nR 0” to retrieve all possible oligos without any overlap. CombinedDensityReduction script provides two files: a Hit archive showing all the *hits* and, a non-redundant list of oligos (BED). The Hit file is used for statistical analysis of oligos along the target scaffold using R software, whereas the non-redundant list of oligos is used for library construction.

### Generation of shared oligopaints between Y3 and other Y-linked scaffolds

Since the density for Y3 scaffold was notably low as 0.09 and 0.12 *hits*/kb for single copy and repetitive oligos respectively, we took advantage of the other 38 Y-linked scaffolds to design shared oligos between Y3 and these other scaffolds. To begin, a shared oligo is, by definition, a repetitive oligo. Thus, a Y3 share oligo is one that appears in at least one additional Y-linked scaffold besides Y3. Furthermore, this oligo could be found at more than one location on the Y3 or other scaffold. Briefly, 41 Y-linked scaffolds were concatenated (Y41c) with 50 Ns inserted between scaffolds to avoid creating false oligos. Similarly, Y1, Y2 and Y3 scaffolds were concatenated (Y123c). Repetitive oligos were independently designed for these two concatenated sequences using the method described in the previous section. Then, repetitive oligos of Y123c were subtracted from Y41c, and oligos between the coordinates of Y3 in the Y41c concatenated sequence were selected. These oligos represent the Y3 share oligos. Next, oligos were filtered including kmerFilter, structureCheck, female short reads filter as done previously. Finally, this list of oligos was processed to remove overlap and identical/redundant oligos using the DensityReduction script with “-n 0” and the bash command “uniq -u” respectively. Thus, the resulting list consisted of 4271 oligos which represents the shared oligos between Y3 and the other 38 Y-linked scaffolds (Y3-share: Y3-sh).

### Probe construction

There are different methods for producing and labeling oligopaints at the laboratory, and depending on this, the final composition of the probe may vary. Because our oligopaints probes were produced using amplification by the T7 enzyme from a complex oligo pool (∼90,000 oligos), the final template molecule for every probe contained three components: i) a central targeting sequence for *in situ* hybridization to the target scaffold/chromosome, i.e. the oligos designed in the previous sections, ii) a flanking priming region for reverse transcription (RT) and, iii) two index flanking priming regions (forward and reverse) to allow selection/amplification of a subset of oligos required for a specific experiment (S13 Fig). The flanking priming regions were adopted from Beliveau et al., (2015) and Chen et al., (2015) (S1 Table). All flanking primers showed absence of similarity among them and with the 5 nt of the T7 promoter’s 3’ terminal region (TAGGG). This was accomplished using BLAST (Altschul S. et al., 1990). The ∼180,000 oligos designed for *D. miranda* were divided into two 90K oligopools and purchased from GenScript. The first oligopool contained oligos for chr4, XR-A and Y, while the second contained oligos for XR-D, XL and neo-X. Flanking primers were screened against every oligopool using BLAST, and the primers with fewer alignments were selected to construct the final probes for that particular oligopool. The complete list of oligos for every chromosome can be found in S2 Table and S3 Table.

### Probe production

The production of the probes was carried out following the protocol described by Nguyen and Joyce (2019). Briefly, the oligo library of the target chromosome was selected and amplified using the index primers through real-time PCR (Applied Biosystems StepOne). A 50 μl PCR reaction was assembled with the following components: 2.5 μl of 10 μM index forward primer, 2.5 μl of 10 μM index reverse primer, 2.5 μl of 1 ng/μl oligo pool, 2.5 μl of 20X EvaGreen, 12.5 μl of ddH2O, and 25 μl of Phusion Hot Start II High-Fidelity PCR Master Mix (Thermo Fisher). The PCR program consisted of an initial denaturation at 98°C for 3 min, followed by 24-36 cycles of denaturation at 98°C for 5 sec, annealing at 72°C for 15 sec, and a final hold at 4°C. A 3% agarose gel electrophoresis in 1X TBE was used to confirm the selective amplification of the libraries, observing a single band. The PCR products were purified using DNA Clean & Concentrator®-5 (Zymo Research) and quantified using Nanodrop Lite.

Subsequently, RNA in vitro transcription of the PCR products was performed using the MEGAscript™ T7 Transcription Kit (Thermo Fisher). A 22 μl reaction was assembled with the following components: 8 μl of NTP mix, 2 μl of 10X reaction buffer, 1 μl of RNaseOut (Invitrogen), 2 μl of T7 polymerase, and 9 μl of the PCR product. The reaction was incubated at 37°C overnight (∼12-14 hr). On the following day, the generated products were fluorescently labeled through reverse transcription using the Maxima H Minus Reverse Transcriptase Kit (Thermo Fisher). Thus, a 60 μl reaction was assembled with the following components: 7 μl of dNTP mix (10 mM each), 14 μl of Maxima H Minus RT buffer, 2 μl of RNaseOut, 1.5 μl of Maxima H Minus RT enzyme, 3.5 μl of ddH2O, 10 μl of RT-fluorophore primer (100 μM), and the 22 μl of the previously generated product. The reaction was incubated at 50°C for 90 min using a thermocycler (Bio-Rad). Then, 50 μl (1:1 v/v of 1M NaOH: 0.5 M EDTA) was added to the reaction tube and incubated at 95°C for 10 min to digest the RNA strand in the oligos labeled with Cy5 and FAM fluorophores. In the case of oligos labeled with ATTO488, alkaline digestion was avoided as the fluorophore is sensitive to the reaction. 240 μl of Oligo binding buffer (Zymo Research) and 960 μl of absolute ethanol were added to the previous reaction. The components were mixed, and finally, the oligos were purified using DCC-25 (Zymo Research). Lastly, the oligos were diluted in ddH2O.

### Oligopaint FISH on slides

Male and female 3rd instar larvae were separated based on testes identification under Leica EZ4 stereomicroscope. Larval brains were dissected in PBS solution at room temperature (RT) in a concave slide. Approximately 10-12 brains were collected and washed twice with PBS. Subsequently, they were hypotonically treated with 100 μl of 0.075 mM KCl for 10 minutes at RT. 30 μl of the hypotonic solution was removed and the brains were first fixed by adding 30 μl of fresh Carnoy’s solution (3:1 v/v of absolute ethanol:glacial acetic acid) to the hypotonic solution for 5 minutes, followed by a second fixation with 100 μl of Carnoy’s solution for 5 minutes. The brains were collected in a microtube containing 100 μl of 60% glacial acetic acid and gently homogenized to obtain a cell suspension. Finally, 25 μl aliquots of the cell suspension were placed on each slide and allowed to dry. The slides were stored at −20°C until the moment of use. Next steps were carried out based on the protocol described by Nguyen and Joyce (2019) with some modifications. Briefly, slides with fixed cells are incubated in 1X PBS for 10 minutes at RT and PBS-T (1X PBS + 0.5% (vol/vol) Triton X-100) for 15 minutes at RT. The slides are dehydrated in 70% ethanol for 2 minutes, 85% ethanol for 2 minutes and 100% ethanol for 2 minutes at RT. Subsequently, the slides are permeabilized with 2X SSCT (0.3 M NaCl, 30 mM sodium citrate, 0.2% (vol/vol) Tween-20) for 5 minutes at RT. The slides are pre-treated in 2X SSCT + 50% formamide for 5 minutes at RT, 2X SSCT + 50% formamide at 92°C for 2 minutes, and 2X SSCT + 50% formamide at 60°C for 15 minutes. The hybridization mix (25 μl) was prepared with the following components: 12.5 μl of formamide, 6.25 μl of dextran sulfate mix (0.4 g/ml dextran sulfate, 8X SSC, and 0.4% (vol/vol) Tween-20), 1 μl of RNAseA (10 mM each), 1.85 μl of dH2O, and 2 μl of the oligopaints library (or 18S rDNA-digoxigenin (*Helicoverpa Zea*) or 21mer-biotin (5’AAAACTTATCTCCGCTGGCGG3’) centromeric probe). We used at least 25 pmol of oligopaints per slide. 25 μl of the hybridization mix was placed per slide and covered with a coverslip. The slides were denatured using a dry bath at 72°C for 2 minutes and then incubated overnight in a humid chamber using 37°C or 47°C for *D. miranda* and 37°C for *D. pseudoobscura*. On the following day, the coverslips were removed and the slides were washed twice in 2X SSC at 42°C for 5 minutes and 0.2X SSC at 42°C for 5 minutes. Finally, the slides were incubated in 2X SSC at RT for 15 minutes. DNA counterstain was performed with 13 μl of Vectashield or SlowFade™ (Invitrogen) with DAPI. Images were acquired in a Zeiss Axioplan 2 microscope equipped with AxioCam MRm and a 100X Plan Apochromat oil immersion objective lens (NA = 1.4) (Carl Zeiss, Inc., Jena, Germany) and Isis V5.2 (MetaSystems GmbH) software. GIMP 2.10.20 (The GIMP Development Team, 2019) and ImageJ (Schneider et al., 2012) were used for image processing. The resolution of the images for the DAPI channel was improved using deconvolution. Then, point-spread function (PFS) were calculated using the Born and Wolf 3D optical model implemented in the plugging PFS generator (Kirshner et al., 2013) for ImageJ. The model was run with settings according to our microscope (1.4 for numerical aperture NA, 67.5 for pixel size XY), 160 nm for Z-step (25 steps) and 461 nm of emission wavelength (DAPI). Next, the point-spread function was used to run deconvolutionLab2 (Sage et al., 2017) using the Richardson-Lucy algorithm with 50 iterations in ImageJ.

### Chromosomal measurement

Chromosomal measurements of the *Drosophila miranda* Y chromosome were performed using 25 independent metaphase microphotographs. Scaffold Y2, which exhibited uniform fluorescence along its entire length, was used as a reference. Its known genomic length (37.2 Mb) served as the calibration standard for measuring distances within the Y chromosome using ImageJ software (S8 Fig, A–III).

### Whole-Chromosome alignments

Large-scale alignments were conducted between chromosome 3 of *D. pseudoobscura* MV25 (GenBank NC_046680.1) and the Y1 and Y2 scaffolds of *D. miranda*. The Nucmer software with parameters "-t 4 -L 2500" was used for initial alignment (Marçais et al., 2018). Subsequently, delta-filter was applied with the "-r -q" options to filter and refine the alignments. Only alignments exceeding 2500 nucleotides were visualized using *genoPlotR* (Guy et al., 2010). To highlight regions of interest, alignments within 1Mb windows from the *fest* gene (11136066 nt) to the centromere were color-coded (Fig 5C).

### Enhancing oligo density estimation

To accurately estimate oligo densities between species and correct for repetitive library biases, oligos were aligned using Bowtie2 with the following parameters: -D 20 -R 3 -N 1 -L15 -i S,1,0.50 -score-min G,30,10 -local. These settings deviate from those used for oligo design, increasing search sensitivity (-L15), which is crucial for Y-linked oligos. Moreover, a stringent alignment criterion (up to 2 mismatches or 1 gap) was adopted by setting “G,30,10” as minimum score threshold. To avoid double-counting, overlapping alignments were removed and oligo densities were then calculated per kilobase (kb) within 100 kb windows. Employing these parameters, we observed a strong correlation between our FISH results and the alignment-based density estimates. Then, we successfully identified oligo alignments within the *fest* gene on chromosome 3 (Fig 5B), and telomeres on the dot chromosome (S8 Fig, B) of *D. pseudoobscura* using the Y3-Sh library of *D. miranda*. Additionally, the Y3-U, Y3-R, and Y3-Sh libraries revealed oligo alignments within the *fest* gene located approximately 17Mb of the Y1 scaffolds (S8 Fig, A) and not alignment on the Y2 as seen in FISH. Finally, it allowed us to refine the density estimations for repetitive libraries, including XR-A, XL-1, Y1, Y2, and Y3, resulting in values that more closely correspond to expected fluorescent detection levels (Table B in S1 Text).

## Acknowledgments

We extend our sincere gratitude to the members of our labs and colleagues for their valuable insights and stimulating discussions. We also acknowledge the financial support provided by the Fundação de Amparo à Pesquisa do Estado de São Paulo (FAPESP) (MDV: 15/20844-4 and 23/14607-6; HBB: 19/10559-1).

## Supporting information

**S1 Fig. Implementation of single-copy and repetitive oligos. A)** Detail pipeline for single-copy oligos based on OligoMiner (Beliveau et al., 2018). **B)** Detailed pipeline for repetitive oligos based on OligoMiner and OligoY (Almeida et al., 2024.). A pipeline to design oligos for the Y1 scaffold (gray rectangle) is presented where larger colored rectangles correspond to distinct scaffolds in the genome, while the shorter rectangles represent oligos. Oligos of different colors indicate distinct sequences. Note that for repetitive oligos, the Y1 scaffold is absent during the Bowtie2 alignment step. This enables the retrieval of both single-copy and repetitive oligos. ^+^Female short read step is added only for Y-linked oligos. *The final filtering step, Density reduction, is applied to obtain single-copy or repetitive oligo but not both. For a combined list of single-copy supplemented with repetitive oligos, Density reduction is skipped, and instead, Combine density reduction is used. B and C conclude generating a BED list of oligos that will be used to probe construction. For further details, refer to the Materials and Methods section.

**S2 Fig. Oligo-FISH on Neo-X and chromosome 4 of *D. miranda*. A)** and **B)** display two male metaphases labeled with single-copy oligos (Cy5: magenta). **C)** and **D)** show two male metaphases targeted with single-copy oligos (Cy5: cyan) and rDNA probe (antiDIG-rhodamine: blue). Bar scale, 2.5 μm.

**S3 Fig. Violin plot of the distance between oligos for the XL/XR chromosome.** The y-axis displays the distance, in nucleotides, computed using a window of 1Mb or 2Mb, and transformed to log10. The horizontal red dash line indicates a distance of 1000 nucleotides. The x-axis shows the chromosome position of each window. The black point inside each violin represents the median and its vertical lines the quantiles 0.25 and 0.75. **A)** Violin plot of single-copy oligos in the scaffold XR-D. **B)** Violin plot of single-copy oligos in the scaffold XL-A. **C)** Violin plot of the scaffold XR-A for (I) the single-copy and (II) repetitive oligos. **D)** Violin plot of the scaffold XL-1 for (I) the single-copy and (II) repetitive oligos.

**S4 Fig. Violin plot of the distance between oligos for A) Neo-X and B) chr4.** The y-axis displays the distance computed using a window of 2Mb and transformed to log10. The red dash line indicates a distance of 1000 nucleotides. The x-axis shows DNA sequence position on each chromosome for each window. The black point inside each violin represents the median and its vertical lines the quantiles 0.25 and 0.75.

**S5 Fig. Oligo-FISH on the XL/XR chromosome of *D. miranda*. A)** and **B)** show two male metaphases labeled with four single-copy and two repetitive oligos libraries along the XL/XR chromosome. I) displays the mapping of the six libraries and DAPI for DNA, while from II) to VII) represent the labeling of every library alone. II: XRD-Sc (Cy5: green), III: XRA-Sc (atto488: red), IV: XRA-R (atto488: blue), V: XL1-Sc (Cy5: cyan), VI: XL1-R (Cy5: yellow), VII: XLA-Sc (atto488: magenta). VIII shows only the labeling of the oligopaint libraries. Note the lack of targeting between XL1 and XLA, corresponding to the rDNA region. Both single-copy and repetitive libraries demonstrate no off-target hybridization. Bar scale, 2.5 μm.

**S6 Fig. Violin plot of the distance between oligos for the Y1 and Y2 scaffolds.** The y-axis displays the distance, in nucleotides, computed using a window of 2Mb, and transformed to log10. The red dash line indicates a distance of 1000 nucleotides. The x-axis shows the DNA sequence position of each window along the scaffolds. The black point inside each violin represents the median and its vertical lines the quantiles 0.25 and 0.75. **A)** Violin plot of (I) single-copy and (II) repetitive oligo libraries for the scaffold Y1. **B)** Violin plot of (I) single-copy and (II) repetitive oligo libraries for the scaffold Y2.

**S7 Fig. Oligo-FISH on Y chromosome of *D. miranda* using repetitive libraries** A), B) and C) are male metaphases. **A)**. FISH using repetitive oligos for Y1 (Cy5: magenta) and Y3 (atto488: green). **B)** FISH using repetitive oligos for Y2 (atto488: magenta) and ribosomal (rDNA) probe (antiDIG-rhodamine: cyan). **C)** FISH using Y3-share oligos (atto488: green) and rDNA probe. A, B and C demonstrate the high specificity of the repetitive oligo libraries, with no off-target hybridization. Sc: single-copy, R: repetitive library, Sh: shared library. Bar scale, 2.5 μm.

**S8 Fig. Enhancing oligo density estimation. A)** Explanation of off-target label of Y3 oligos (B label in panel A.III) in Y1 scaffold using Bowtie alignment (see Enhancing oligo density estimation in Material and method section). The method was validated aligning the libraries from scaffold Y3 on scaffold Y2, resulting in minimal density values along it (panel A.II), consistent with the absence of labeling in FISH experiments (panel A.III). Panels A.I and A.II represent the density (no. *hits*/kb) of single-copy (Sc), repetitive (R) oligos from Y3 along the Y1 and Y2 scaffolds respectively. The x-axis shows chromosome position (Mb). Notably, there is a heightened density around the 17 Mb position on Y1, which accounts for the observed B interstitial labeling on this chromosome. This is consistent with the chromosomal measurements (using 25 independent metaphases) between the start of Y1 (α) and the label B (β), as indicated by the orange line in panel A.III, which measures 17.15 ± 2.1 Mb. A.III. Chromosome Y targeted with Y3-shared (atto488: green) and centromeric 21mer (Avidin-FITC: yellow) probe. **B)** Estimating oligo densities between species. The Y3-Sh library of *D. miranda* aligned at the end of the Dot chromosome of *D. pseudoobscura* (*D. pse*) revealing sufficient density of oligos (near 0.25 *hits*/kb) to be detected by FISH. Two male metaphases subjected to FISH using Y3-Sh library in *D. pse* demonstrating label in the telomere of the Dot chromosome (orange heat arrow). Bar scale, 2.5 μm.

**S9 Fig. Oligo-FISH in *D. pseudoobscura* using oligo libraries of *D. miranda*. A)** FISH using single-copy oligos of Y1 (atto488: blue) and Y2 (cy5: cyan), targeting the region medial to proximal to the centromere, and medial to the telomere of the Muller C (chr 3), respectively. **B)** FISH with single-copy oligos of Y3 (Y3-Sc) (cy5: green) labeling all the YD. **C)** FISH using repetitive oligos of Y3 (Y3-R) (atto488: green) showing three blocks of hybridization along the YD. **D)** FISH with Y3 shared oligo (Y3-share) (atto488: orange) demonstrating hybridization in telomeric regions of autosomes and YD (arrows). Furthermore, there is a label in the chr3 which represents the FBgn0082139 gene (ortholog to *fest* gene of *D. melanogaster*). **E)** FISH using Y1-Sc (cy5: blue) and Y3-share (orange) oligos. Note that the interstitial label is located at the end of the Y1 label on chr3 (arrow). **F)** FISH using Y2-Sc (cy5: cyan) and Y3-share (orange) oligos revealed that Y2 labeling extends from the middle to the end of the chr3 (Muller C). Panels E and F illustrated that the FBgn0082139 gene lies precisely between the Y1 and Y2 labels on chr3. All panels are male metaphases of *D. pseudoobscura*. Bar scale, 2.5 μm.

**S10 Fig. Oligo-FISH using *fest* and *pain* oligo libraries in *D. miranda*. A, B and C)** FISH using single-copy oligos for *fest* gene (atto488: green) targeting an internal region in Y1 (Label B in Fig4B, V) and inside Y3 (Label C in Fig4B, V). **D, E and F)** FISH using single-copy oligos for *pain* gene (atto488: red) labeling the end of the left arm (Label A in Fig4B, V) and inside Y3.

**S11 Fig. Alternative scenario for the evolution of the *D. miranda* Y chromosome.** A potential evolutionary pathway for the *D. miranda* Y chromosome involves a translocation between the long arm of the YD chromosome and the long arm of the Müller C chromosome. This translocation would have created the interstitial telomere observed in *D. miranda*. However, the resulting chromosome would have been acrocentric, differing from the observed metacentric structure. To explain the formation of the discontinuous Y1 and Y2 scaffolds, a paracentric inversion is hypothesized. Yet, this inversion would have repositioned the *fest* gene to the end of Y2 and placed Y2 within the YD regions (green), which contradicts the FISH experimental data

**S12 Fig. Identification of Y-linked scaffolds. A)** YGS analysis using 18-mers. The graph shows the percentage of unmatched valid single-copy (VSC) female k-mers in the scaffolds and the size of the scaffolds (log10kb). The vertical line indicates the minimum threshold (80%) to consider a Y-linked scaffold. The scaffolds are shown according to Mahajan et al. (2018) as follows: A (autosome), A/X (autosome or X chromosome), Y (Y-linked). **B)** Intersection of the three lists of classifications (YGS 15-mers, YGS 18-mers and coverage male/female reads) to identify Y-linked scaffolds. A total of 41 scaffolds were assigned as linked to the Y chromosome, including the three largest scaffolds Y1, Y2, Y3, and 38 other shorter scaffolds.

**S13 Fig. Composition of the oligo sequence. A)** An oligo probe consists of the target genomic region that will be hybridized in the FISH experiment and priming regions used during probe production. **B)** Complex oligo library. The forward and reverse index priming regions are unique to each sub-library within the oligo pool. Therefore, it is possible to select sub-libraries during the oligo production in the laboratory.

**Supplemental tables. S1 Table.** List of the flanking priming regions used to construct the oligo libraries. **S2 Table.** Oligos for Muller-B and Xs chromosomes. **S3 Table.** Oligos for the chromosome Y.

**Supplemental text. Table A. Number of oligos, no. de *hits* and density (no. *hits*/kb) for each target scaffold.** The spacing (-n: nucleotides) between oligos used in DensityReduction.py step is shown for every library. Mb: Megabases. **Table B. Enhancing oligo density estimation.** Bowtie alignment (see Enhancing oligo density estimation in Material and method section) revealed a notable increase in densities for repetitive (R) oligo libraries while single-copy (Sc) oligo libraries remained similar. Exact alignments represent identical oligos to the target sequence, while relaxed alignments allow for up to 2 mismatches or 1 gap. Density (no. *hits*/kb). **Table C.** Density estimation of single-copy (Sc) and repetitive (R) oligos libraries for Y3 in other Y-linked scaffolds using the Enhancing oligo density estimation (see Material and method section). Density (no. *hits*/kb).

